# Contributions of different prefrontal cortical regions to abstract rule acquisition and reversal in monkeys

**DOI:** 10.1101/180893

**Authors:** Giancarlo La Camera, Sebastien Bouret, Barry J. Richmond

## Abstract

The ability to learn and follow abstract rules relies on intact prefrontal regions including the lateral prefrontal cortex (LPFC) and the orbitofrontal cortex (OFC). Here, we investigate the specific roles of these brain regions in learning rules that depend critically on the formation of abstract concepts as opposed to simpler input-output associations. To this aim, we tested monkeys with bilateral removals of either LPFC or OFC on a rapidly learned task requiring the formation of the abstract concept of same vs. different. While monkeys with OFC removals were significantly slower than controls at both acquiring and reversing the concept-based rule, monkeys with LPFC removals were not impaired in acquiring the task, but were significantly slower at rule reversal. Neither group was impaired in the acquisition or reversal of a delayed visual cue-outcome association task without a concept-based rule. These results suggest that OFC is essential for the implementation of a concept-based rule, whereas LPFC seems essential for its modification once established.

## Introduction

Primate prefrontal cortex (PFC) supports executive, mnemonic and attentional functions critical for learning and invoking rule-based strategies to control behavior (Miller and Cohen, 2001; Curtis and D'Esposito, 2004; Bunge et al., 2005; Tanji et al., 2007; Fuster, 2008; Tanji and Hoshi, 2008; Wise, 2008; Buckley et al., 2009). Different prefrontal regions seem to support different aspects of rule-based behavior. Lateral prefrontal cortex (LPFC) has been implicated in working memory (Fuster and Alexander, 1971; Funahashi et al., 1989, 1993), attention (Lebedev et al., 2004), executive control (Huettel et al., 2004; Tanji and Hoshi, 2008), self-organized behavior (Procyk and Goldman-Rakic, 2006), rule-based behavior (Rushworth et al., 1997; Wallis et al., 2001; Bunge et al., 2003; Shima et al., 2007; Tanji et al., 2007; Moore et al., 2009, 2012), and context-dependent decisions (Wise et al., 1996; Mante et al., 2013; Rigotti et al., 2013), whereas orbitofrontal cortex (OFC) has been implicated in reversal learning (Dias et al., 1996, 1997; Izquierdo et al., 2004; Walton et al., 2010), reinforcement learning (Rolls et al., 1996; Rolls, 2000; Hampton et al., 2006; Salzman et al., 2007; Simmons and Richmond, 2008; McDannald et al., 2011), reward evaluation and comparison (Tremblay and Schultz, 1999; Schultz et al., 2000; Wallis and Miller, 2003; Salzman et al., 2007; Bouret and Richmond, 2010; Simmons et al., 2010), and the evaluation of alternative options (Padoa-Schioppa and Assad, 2006, 2008).

In monkeys, OFC seems to have a role in decisions based on expected outcome value beyond simple stimulus-response associations (Walton et al., 2010; Clark et al., 2013), whereas LPFC seems essential in updating a rule-based strategy to optimize a rewarding outcome (Dias et al., 1996; Buckley et al., 2009; Moore et al., 2012). An important determinant of reward-based learning is the nature of the predictive cues used for learning, which may be simple stimuli acting in isolation, compound stimuli, or abstract concepts. Presumably, a concept-based task such as a DMS task (“if the 2 stimuli match, then reward”) results in different cognitive demands than a simpler cue-outcome association task (“if stimulus A, then reward”). Since LPFC is implicated in rule learning and OFC is implicated in assessing outcome value, we tested the effects of LPFC and OFC removals in learning a behavior that requires the formation of abstract concepts compared to a behavior that requires simple visual stimulus-outcome associations. Normal monkeys and monkeys with bilateral LPFC and OFC lesions were tested in a DMS task (requiring the formation of the concepts of ‘same’ and ‘different’) and in two simpler rule-based tasks that required no concept formation. This comparison should expose significant differences in the roles of LPFC and OFC in behavior that depends on concept-based as opposed to sensory-cue based predictions of forthcoming contingencies.

To be able to compare the learning times across all tasks we developed a variation of DMS that was learned rapidly by the monkeys. Control monkeys quickly learned to use the association between the abstract concepts of ‘same’ and ‘different’ with their predicted outcome. Monkeys with bilateral OFC lesions were impaired at both acquiring and reversing the associations between the concept and the outcome, whereas monkeys with bilateral LPFC lesions acquired the task as quickly as the control group, but were impaired at reversing the association between concept and outcome. Both lesion groups learned the simpler cue-outcome associations (with and without a memory component) as quickly as controls, showing that the impairments were related to forming the abstract concept and/or applying the abstract concept to infer the rule of the task. These results suggest that OFC is essential for acquiring and updating an association between a concept and a reward, whereas LPFC seems essential for its modification once established.

## Methods

### Subjects and surgical procedures

Nine rhesus monkeys were used for this study, 3 unoperated controls, 3 monkeys with bilateral LPFC lesions and 3 monkeys with bilateral OFC lesions (see below for details). All the experimental procedures were carried out in accordance with the ILAR Guide for Care and Use of Laboratory Animals and approved by the Animal Care and Use Committee of the National Institute of Mental Health. Monkeys received bilateral lesions of orbital or lateral prefrontal cortex using a combination of suction and electrocautery. The intended lateral prefrontal lesion (Figure 1A) extended laterally from the dorsal midline to the orbital surface of the inferior convexity. The rostral limit of the lesion was the frontal pole. The caudal limit was the caudal extent of the principal sulcus. The frontal eye fields and the banks of the arcuate sulci were intentionally spared. In total, the intended lateral prefrontal lesion included areas 9, 46, 45, 12, and dorsal area 10 (Walker, 1940; Petrides and Pandya, 1994). The intended orbital prefrontal lesion (Figure 1B) extended from the fundus of the lateral orbital sulcus to the fundus of the rostral sulcus. The rostral limit of the lesion was a line joining the anterior tips of the lateral and medial orbital sulci. The caudal limit was approximately 5 mm rostral to the junction of the frontal and temporal lobes. In total, the intended orbital prefrontal lesion included areas 11, 13, 14 and the caudal part of ventral area 10 (Walker, 1940; Petrides and Pandya, 1994). The lateral and orbital prefrontal lesions shared a common boundary at the lateral orbital sulcus.

**Figure 1:**
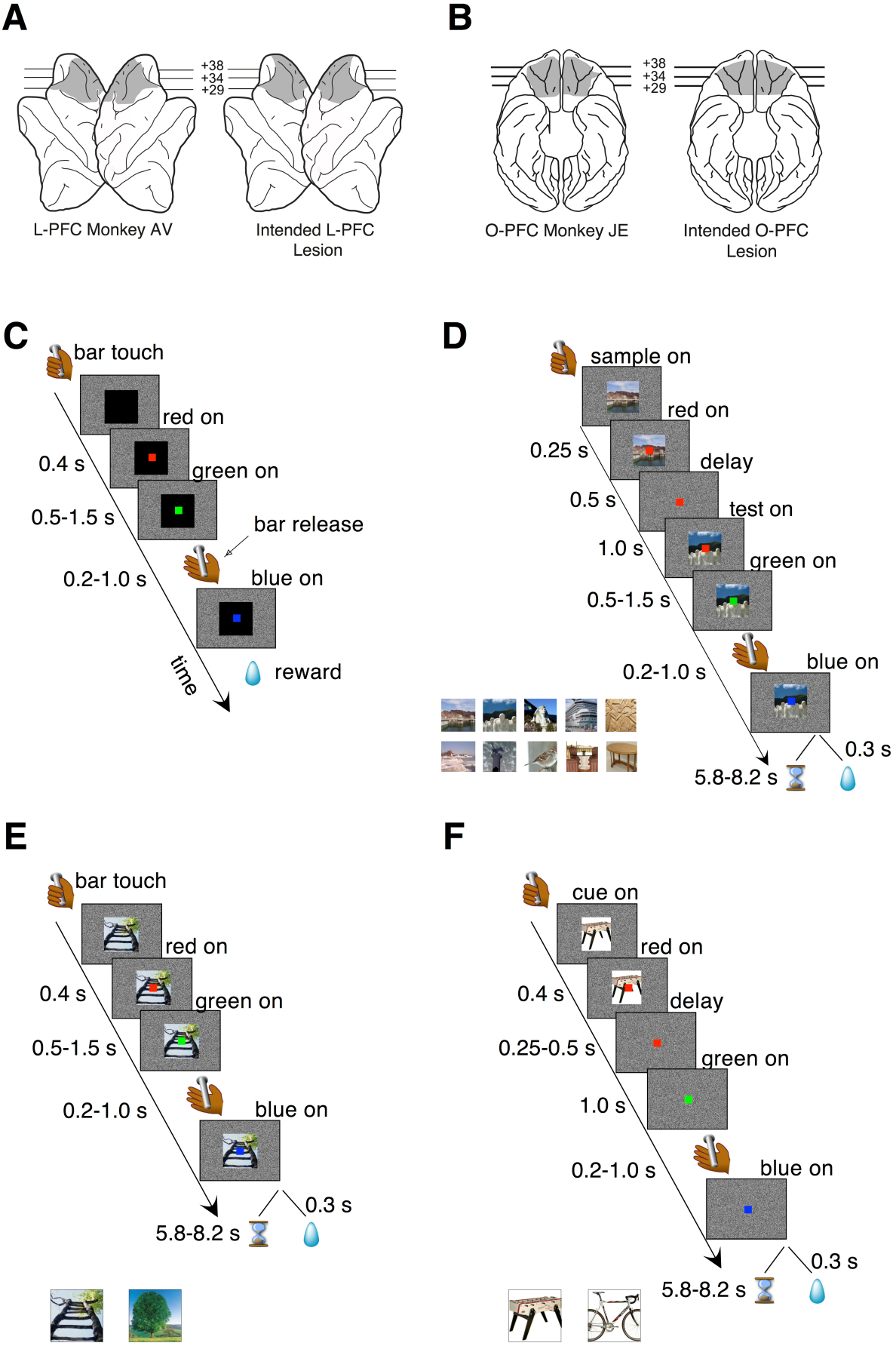
Intended lesions and behavioral tasks. **A)** Lateral view of a standard rhesus monkey brain showing the extent of the LPFC lesion in monkey AV (left, shaded region) and the intended lateral PFC lesion (right, shaded region). **B)** Ventral view of a standard rhesus monkey brain showing the extent of the OFC lesion in monkey JE (left, shaded region) and the intended orbital PFC lesion (right, shaded region). **C)** Bar release task. The monkeys were required to release a bar when a red dot, located centrally on a computer screen, turned green. Failure to do so resulted in an incorrect trial that was immediately aborted. Correct trials were rewarded with a liquid reward; incorrect trials were aborted and no reward was given. Transition times between events are reported to the left (see Methods for details). **D)** DMS task. Two visual cues, a ‘sample’ and a ‘test’ image, were presented in sequence and separated by a temporal delay of 1 second. The sample image was turned off after 500ms from its onset, starting the delay period during which only the red dot was visible on the screen. At the end of the delay period, the test image appeared on the screen behind the red dot; between 500 and 1500ms later, the red dot turned green. A bar release at this point (between 200ms and 1000ms after the onset of the green dot), led to either reward or a time-out depending on the rule in effect. A bar release outside the allowed interval caused the immediate abortion of the trial (see Methods for details). *Inset* at the bottom shows a sample of the cue set. Sets of 50 images were used in each session. **E)** Simple association task where one of two visual cues was associated to reward while the other predicted a time-out. Regardless of the outcome of the previous trial, each cue had an equal chance of being selected in the following trial. Transition times between events are reported to the left and the two visual stimuli used as cues are shown below the main plot. All other task features were as in the DMS task of Figure 1D (see Methods for details). **F)** Delayed association task. In this task, the visual cue disappeared after 250-500ms from the onset of the red dot, which turned green after an additional delay period of 1 second. Everything else was as in the non-delayed association task of panel E. Two different visual cues (shown below the main plot) were used. Panels A and B adapted from (Simmons et al., 2010).

Lesion location and extent were largely as intended within each experimental group. In the LPFC group, all three lesions extended from 23 to 43 mm rostral to the interaural line. All LPFC lesions included regions 9, 46, 45, and 10, as intended. However, most of 12o (on the orbital surface) and the caudal part of 12l (on the ventrolateral surface) were spared in all three animals. The banks of the arcuate sulcus were spared, as intended. There were no areas of unintended damage. In the OFC group, two of the three lesions (JE and SU) extended from 25 to 42 mm rostral to the interaural line; the other (BR) was placed slightly more caudally, extending from 23 to 41 mm rostral to the interaural line. Except for a narrow strip of area 14 immediately ventral to the rostral sulcus and a small part of area 13l immediately medial to the lateral orbital sulcus, OFC lesions in all three monkeys included all intended regions. There were no areas of unintended damage. More details are reported in (Simmons et al., 2010).

### Behavioral training

The monkeys were initially trained on a simple bar release task (Figure 1C). They sat in a monkey chair facing a computer screen displaying a white noise background and had access to a bar located on the primate chair. When the monkey touched the bar, a visual stimulus (cue) appeared in the center of the screen (13° on a side; black square in Figure 1C), followed 400 ms later by the appearance of a red dot (0.5° on a side), superimposed to the visual cue in the center of the cue. After a randomly selected time interval of 500, 750, 1000, 1250 or 1500ms, the red dot turned green. The monkey was required to release the bar 200-1000ms from the appearance of the green dot. This caused the green dot to turn blue, followed by the reward (one drop of juice) after 300ms. A bar release outside of the 200-1000ms interval (or not occurring at all within 5 s of the onset of the green dot) caused the trial to be immediately aborted, with all stimuli disappearing from the screen until the next trial. No reward was given after an aborted trial. After each trial, whether correct or not, there was an inter-trial interval (ITI) of 1 second before the monkey could initiate a new trial.

Once the monkeys got proficient in this task (>75% correct), which occurred within 2 weeks for all the monkeys, they were tested to a variation of this task where in some specific correct trials the reward was given only after a delay (‘reward postponement’ task; (Minamimoto et al., 2009). Specifically, three different visual cues (instead of one) were associated with three different reward postponements of 0.3±0.07, 3.6±0.4, and 7.2±0.8 seconds, but the rewards were all equal in size (1 drop of juice). All monkeys (lesioned and controls) learned quickly the meaning of the visual cues (1-3 sessions, *χ*^2^ test for proportions, p<0.01) and developed a linear dependence of error rates vs. predicted reward postponement, following the same pattern reported elsewhere (see Figure 3 of (Minamimoto et al., 2009) and Figure 5 of (Simmons et al., 2010). This test was conducted merely to establish sensitivity to reward postponement, which is instrumental to the main DMS task described next.

### DMS task

In the DMS task (Figure 1D), two visual images were presented sequentially in each trial, initially separated by a temporal delay of 1s. The first image (‘sample’) disappeared after 500ms and only the red dot remained visible. Then the second image (‘test’) appeared and remained on. After 500-1500ms the red dot turned green. In ‘match’ trials, the test matched the sample; in ‘non-match’ trials, the sample and the test cue were different. The stimuli used as sample and test images were chosen randomly from a set of 50 stimuli (see below and inset of Figure 1D for a sample). The monkeys were still required to release the bar on green to complete a trial. The outcome of a completed trial depended on the rule in effect: in the ‘reward-if-match’ rule, a reward would follow if the two images were identical (‘match’ trial), whereas a ‘time-out’ would follow if the two images were different (‘non-match’ trial). In a completed reward trial, the green dot would turn blue and the test cue would disappear, followed by a reward after 300ms, and then by the ITI. In a completed time-out trial, the green dot would turn blue and the test cue would disappear, followed by a pseudo-randomly chosen delay of 5.8-8.2 seconds preceding the ITI. No reward was given in a time-out trial. The opposite would occur in the ‘time-out-if-match’ rule (i.e., time-out would be the outcome of a completed match trial and reward of a completed non-match trial). As in the previous tasks, an incorrect bar release would immediately abort the trial, causing all stimuli to be turned off immediately before entering the 1s ITI. The next trial was always chosen to be a match or a non-match trial with 50% chance, regardless of the behavior in the current trial. The rationale for this task structure was to take advantage of the spontaneous tendency of the monkeys not to release on green when this predicts a time-out (Minamimoto et al., 2009). The monkeys were allowed to engage in the task until they stopped by themselves (they completed an average of 353 ± 103 trials in each session (mean ± SD) prior to the acquisition of the task).

In scoring the monkey’s decisions in the DMS task, we considered ‘correct’ trials either completed trials that led to reward, or aborted trials that would result in a time-out; ‘incorrect’ trials were either completed trials that led to a time-out, or aborted trials that would lead to a reward. Trials with pre-test bar releases (bar releases occurring before the appearance of the test image) were not informative of trial type and were not scored. Bar releases occurring erroneously after the appearance of the test image led to the abortion of the current trial and, in time-out trials, to the avoidance of the time-out. Monkeys learned to do this on purpose and for this reason we termed those trials ‘skipped’ trials. The difference in skip rates between match and non-match trials is an alternative measure of performance, and was used as a criterion to reverse the task’s rule during the experiments (see below). However, since this criterion turned out to be a less reliable measure of performance compared to the correct rates metric defined above, the criterion to task acquisition was defined in terms of percent correct.

The rule of the task was reversed when the difference in skip rates in match vs. non-match trials was significantly different in at least 4 out of 5 consecutive sessions (*χ*^2^ test for proportions, p<0.01). In terms of percent correct, this criterion translated into 2 to 5 consecutive sessions with correct rates significantly above chance (*χ*^2^ test for proportions, p<0.01; see filled circles in Figure 3A). Although less reliable than correct rates, 5 sessions with 4 significant skip rates indicate that the monkey has learned the task, and this criterion insured that all monkeys were asked to reverse the task after similar post-learning experience. Reversal of the task amounted to swapping the association of trial type with reward contingency. For example, match trials, if initially associated with reward, would become associated with time-out, and vice-versa. Rule changes only occurred at the beginning of a new session.

After 5 sessions with a significant separation of skip rates in the reversed task, the delay between sample and test was increased gradually from 1 second to 21 seconds. Duration increments were 1 second per session; only one delay duration was used each day. A delay increment was introduced after each day of significant separation of correct rates with the current delay duration. In most cases, a significant separation of correct rates with new delay duration was reached within the same day the new delay duration was introduced. During testing with longer delays, on occasion (once every seven sessions on average) the delay was kept fixed and a new cue set, never shown before to the monkey, was used. After significant separation of correct rates (which occurred always on the first day a new cue set was introduced), the cue set was kept fixed and the delay incremented by one second, resuming the incremental delay schedule. Thus, once a new set had been introduced, it would be used for a subsequent number of daily sessions until a new set was introduced (or until testing stopped).

We used overall 5 sets of 50 stimuli as sample and test cues. Each visual cue was a 200x200 pixel resolution image. The 50 images in each set represented a large variety of subjects including landscapes, animals, vegetables, man-made objects (planes, cars, tools, etc.; see Figure 1D for a few examples). Different cue sets differed in both the object displayed within a category (e.g., different landscapes in different sets) and the category of objects (e.g., landscapes in one set and tables in another set). Many categories were used in each cue set. A match between sample and test required the stimuli to be identical (e.g., two different images containing similar, but not identical, tables were a non-match).

### Control tasks

After successfully reaching the session with 21 seconds delay, all monkeys were tested in a number of control tasks, described below in the order in which they were executed.

#### - Cue-outcome association task

(Figure 1E). A single visual cue was presented in each trial. There were only 2 cues in this cue set, one predicting reward and the other predicting a time-out. The cue remained on the screen throughout the trial. A bar release after green was required to complete the trial, which would result in reward or time-out depending on which of the two visual cues had been presented. The relative timings of events (except the presentation of the delay and the second image) were as in the DMS task. The stimuli, shown in Figure 1E, had not been used before, but were of same type and resolution as the stimuli used in the DMS task. After the correct rates for the two stimuli had been significantly different for 4 consecutive sessions (*χ*^2^ test, p<0.05), the task was reversed, so that the visual cue previously predicting reward now predicted time-out, and vice-versa.

#### - Delayed cue-outcome association task

(Figure 1F). The monkeys were then tested in a delayed version of the previous task. In this version, the visual cue disappeared between 250 and 500ms after the onset of the red dot, and a 1-second delay followed before the appearance of the green dot. Everything else was as in the non-delayed cue-outcome association task. Two new stimuli were used (Figure 1F). After 4 consecutive days of different correct rates (*χ*^2^ test, p<0.05), the cue-outcome rule was reversed.

#### - Delayed matching-to-sample with short delay

Finally, the monkeys were tested in the DMS task with a sample-test delay interval of 100ms. A 100ms delay was preferred to no delay at all to prevent the monkeys from approaching the task as a perceptual change detection task. The same cue set and abstract rule learned by the monkeys when last exposed to the DMS task were used; percent correct difference was immediately above chance for all monkeys. The monkeys were tested for 4 sessions in this task, then the association between trial type and outcome was reversed. After 4 consecutive days of different correct rates in the two trial types (*χ*^2^ test, p<0.05), the delay between sample and test cue was brought back to 1 second, and a new cue set was used. However, the rule in effect remained the same, and performance (difference in percent correct) was immediately above chance for all monkeys. After 4 sessions, the task was reversed again, and then again 3 more times, each time after 4 consecutive days of significant difference in percent correct (*χ*^2^ test, p<0.05). During these additional reversals, the cue set was not changed.

### Data analysis

All data were analyzed in the R statistical computing environment (R Development Core Team, 2008). Completed trials that led to reward, or aborted trials that would result in a time-out, were scored as ‘correct’; completed trials that led to a time-out, or aborted trials that would lead to a reward, were scored as ‘incorrect’. In the DMS task of Figure 1D, trials with pre-test bar releases (bar releases occurring before the appearance of the test image) were not informative of trial type and were not scored (the monkeys’ behavior could not be the consequence of the predicted outcome).

Performance was quantified as overall percent correct (e.g. (Miyashita, 1988; Miyashita and Chang, 1988; Miller et al., 1993; Amit et al., 1997; Yakovlev et al., 1998; Wallis and Miller, 2003)). In completed trials, reaction times were defined as the time interval between the onset of the green dot and the onset of bar release. Inferences based on reaction times did not alter the conclusions based on correct-rates alone, and thus are not reported.

The first session with percent correct significantly above chance (at p<0.01, *χ*^2^ test), followed by at least one more significant session among the next two, was taken as the onset of discriminative behavior (i.e., as evidence that the association between trial type and reward contingency had been acquired). 99% confidence intervals for percent correct were based on Wilson ‘score’ interval for a binomial proportion (Brown et al., 2001; La Camera and Richmond, 2008).

#### Analysis of the DMS task

After all monkeys had accomplished both acquisition and reversal of the DMS task with one second delay, the number of sessions to criterion were analyzed with a 2-way, mixed design ANOVA, with two factors (‘group’ and ‘protocol’) and within-subjects repeated measures. Post-hoc multiple comparisons were based on a Mann-Whitney test. A Kruskal-Wallis test was used to analyze the number of sessions to criterion for reversal in the DMS task with 100 ms delay.

#### Analysis of control tasks: change-point procedure

For the control tasks of Figure 1E-F we also performed a trials-to-criterion analysis. Trials-to-criterion were obtained with the change-point procedure (Gallistel et al., 2001; Gallistel et al., 2004). The procedure selected a sequence of trials that marked the putative onset of acquisition (or reversal), These trials are called ‘change-points’ and are identified as the points where the change in slope of the cumulative record of correct responses exceeded a chosen criterion. This algorithm follows an iterative procedure: starting from the initial trial *x* and any point *y>x* in the cumulative record, the putative change-point *z* was the point between *x* and *y* placed at the maximal distance from the straight-line connecting *x* and *y*. The selected point was then checked for statistical significance, by comparing performance (percent correct) between trials *x* and *z* vs. performance between trials *z* and *y* (*χ*^2^ test with P-value ‘p’). The putative change-point was accepted as valid if *logit*(p)=log_10_((1-p)/p) was greater than 10 (results were robust to variations in this criterion). Given a valid change-point *z*, the search for the next change-point would start again, this time starting from *x=z* as the initial trial. Otherwise, the procedure was repeated using *y+*1 as the new end point, until a valid change-point (if any) was found. The three following scenarios could occur:

i. At least one change-point was found, in which case the *earliest* change-point was taken as the acquisition/reversal point (Figure 2A);
ii. No change-point was found, but the overall percent correct in the session was significantly higher than chance (p<0.01, *χ*^2^ test). In such a case, the first of 10 consecutive correct responses was taken as the valid change-point (such a point could always be found). This scenario occurred when the cumulative response curve was a straight line, i.e., the integrated correct response rate did not change over the measurement time (Figure 2B);
iii. A special case of ii) occurs when the monkey is indifferent to trial type and always releases on green (the percent correct in this case is 50%). This could happen e.g. when monkeys were first exposed to a task, or in the first session after a task reversal. In those cases, all subsequent sessions were analyzed until a valid change-point was found (typically, this required the analysis of 1-2 additional sessions). All trials in previous sessions were added to the count of the trials performed up until the acquisition/reversal point. This circumstance occurred only infrequently and across all groups.

**Figure 2:**
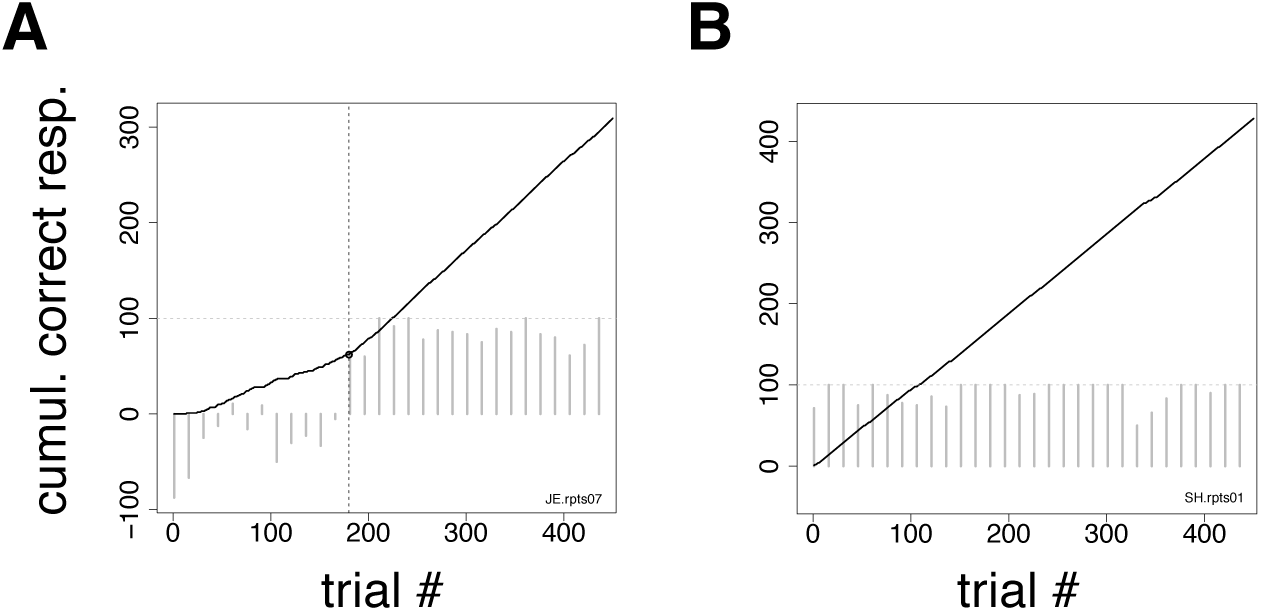
Change-point procedure. Two examples of the change-point procedure used to determine the number of trials-to-criterion in the tasks of Figures 1E-F (see Methods). In both panels, full lines show the cumulative record of correct responses, which grows by one for each correct trial (see Methods). Grey bars show the difference in correct rates between trial types in successive blocks of 15 trials (the horizontal dotted line marks 100% difference in correct rates between trial types). The time-course of the difference in correct rates confirms the validity of the procedure but was not used to find the change-point. **A)** In this example, the change-point (vertical broken line) marks the successful reversal of the cue-outcome association task of Figure 1E on the first day the experimenter had reversed the task (monkey JE). The change-point was reached at trial 180 (corresponding to 48 completed trials). **B)** Cumulative record of correct responses for monkey SH during the first day of exposure to the same task as in panel A. The cumulative record is a straight line implying immediate acquisition of the task, as is apparent from the time course of the difference in correct rates (grey bars). The overall percent correct (with 99% confidence interval) was 91.8 < 95.1 < 97.1. No valid change-point was found in this case, and the first of 10 consecutive correct responses was used as a criterion, according to which acquisition of the task occurred after 6 completed trials.

Since the monkeys received feedback about the rules of the task only in completed trials, to compute the number of trials to criterion we always used the number of completed trials (and not the number of total trials), which were reported as trials-to-criterion. Trials-to-criterion across the groups were analyzed with the same 2-way, mixed design, ANOVA used to analyze the number of sessions to criterion in the DMS task.

## Results

Nine rhesus monkeys, 3 normal controls, 3 with large lateral prefrontal (LPFC) cortex lesions (Figure 1A), and 3 with large orbitofrontal (OFC) lesions (Figure 1B), were initially trained to release a bar when a red dot turned green (Figure 1C). Correct bar releases were rewarded while incorrect ones resulted in the abortion of the trial. After reaching proficiency in this task and after testing all monkeys for sensitivity to reward postponement (Methods), the monkeys were tested in the DMS task of Figure 1D. In this task, they had to predict the outcomes of individual trials (a reward or a 7s ‘time-out’, respectively) based on whether two sequentially presented visual stimuli were the same (‘match’) or different (‘non-match’). Initially, match trials predicted reward and non-match trials predicted a time-out (‘reward-if-match’ rule). As before, a bar release on green was required to complete each trial, which otherwise was aborted; however, a completed trial would now result in a 7s time-out (followed by no reward) in 50% of the trials. The monkeys could learn to ‘skip’ time-out trials by not releasing on green in those trials, which required that the monkeys had learned to infer correctly the trial type according to whether a ‘reward-if-same’ rule or a ‘time-out-if-same’ rule was in effect. In scoring the monkey’s decisions, we considered ‘correct’ either completed trials that led to reward, or aborted trials that would result in a time-out; ‘incorrect’ trials were either completed trials that led to a time-out, or aborted trials that would lead to a reward. Error trials due to a bar release prior to the occurrence of the test image could not be attributed to a predicted outcome and were not scored.

Initially, the overall percent correct was about 50% for each of the 9 monkeys (Figure 3A). This score originated from their previous testing in the bar release task of Figure 1C, when the monkeys had learned to always release on green to obtain reward. After 6-9 sessions, the percent correct for the control and LPFC monkeys increased so that it was greater than chance (Figure 3A, solid black circles, binomial test for proportions, p<0.01). The monkeys with OFC lesions took significantly longer (13-21 sessions) to learn to perform the task to criterion (Figures 3 and 4).

**Figure 3:**
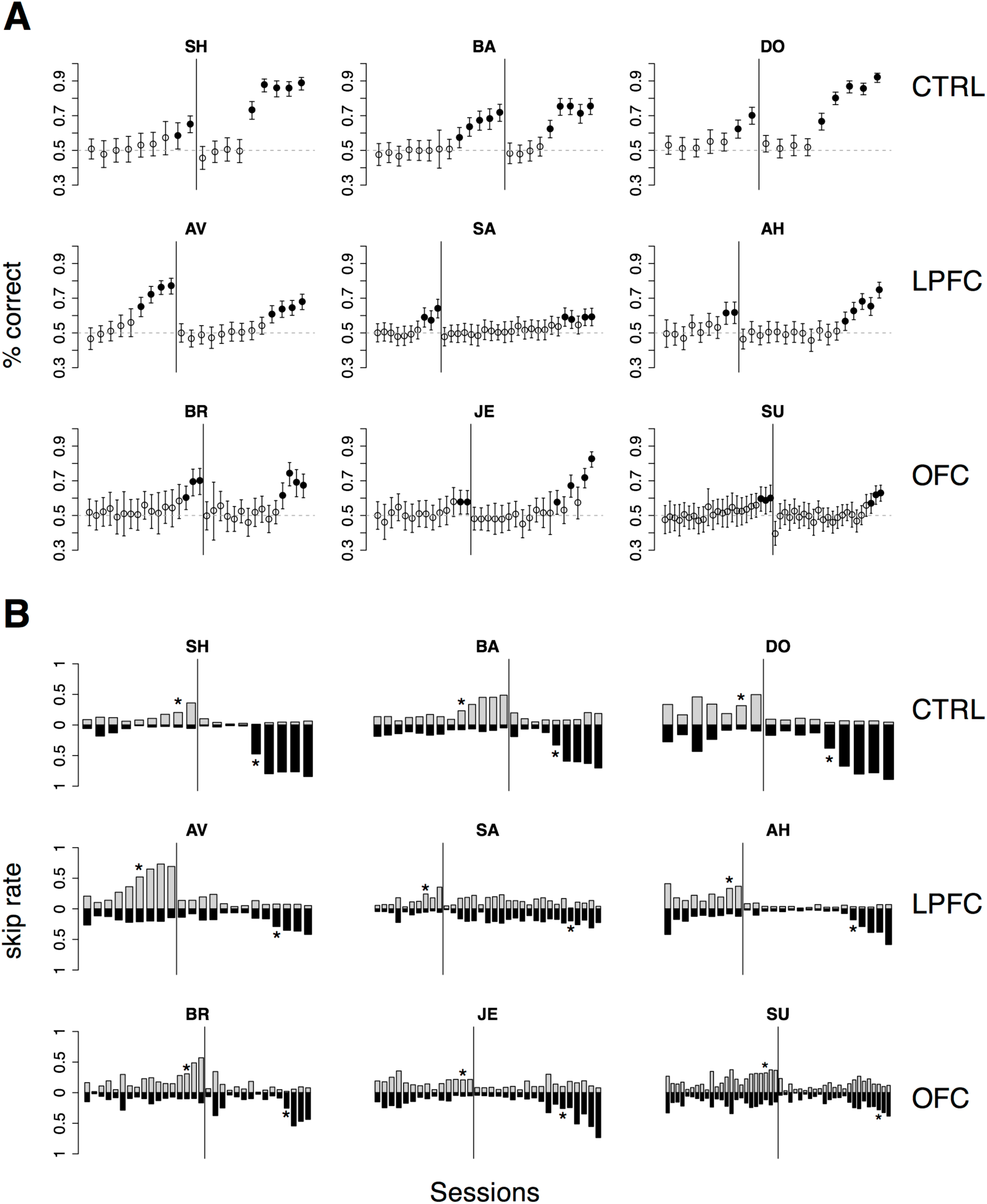
Monkeys’ behavior in the DMS task. **A)** Percent correct vs. session number of all monkeys. Top, middle and bottom rows display data from control (CTRL), LPFC and OFC monkeys, respectively. In each panel, percent correct with 99% confidence intervals are shown (see Methods). Filled circles mark sessions with percent correct significantly above 0.5 (dashed lines; p<0.01, χ^2^ test). Reversal of the task occurred in correspondence of the vertical lines. **B) “**Skip rates” during match (dark bars) vs. non-match trials (lighter bars) in the same data presented in A). The asterisks mark the first day of statistically different % correct performance (as defined in panel A) before (left to the vertical line) and after task reversal.

**Figure 4:**
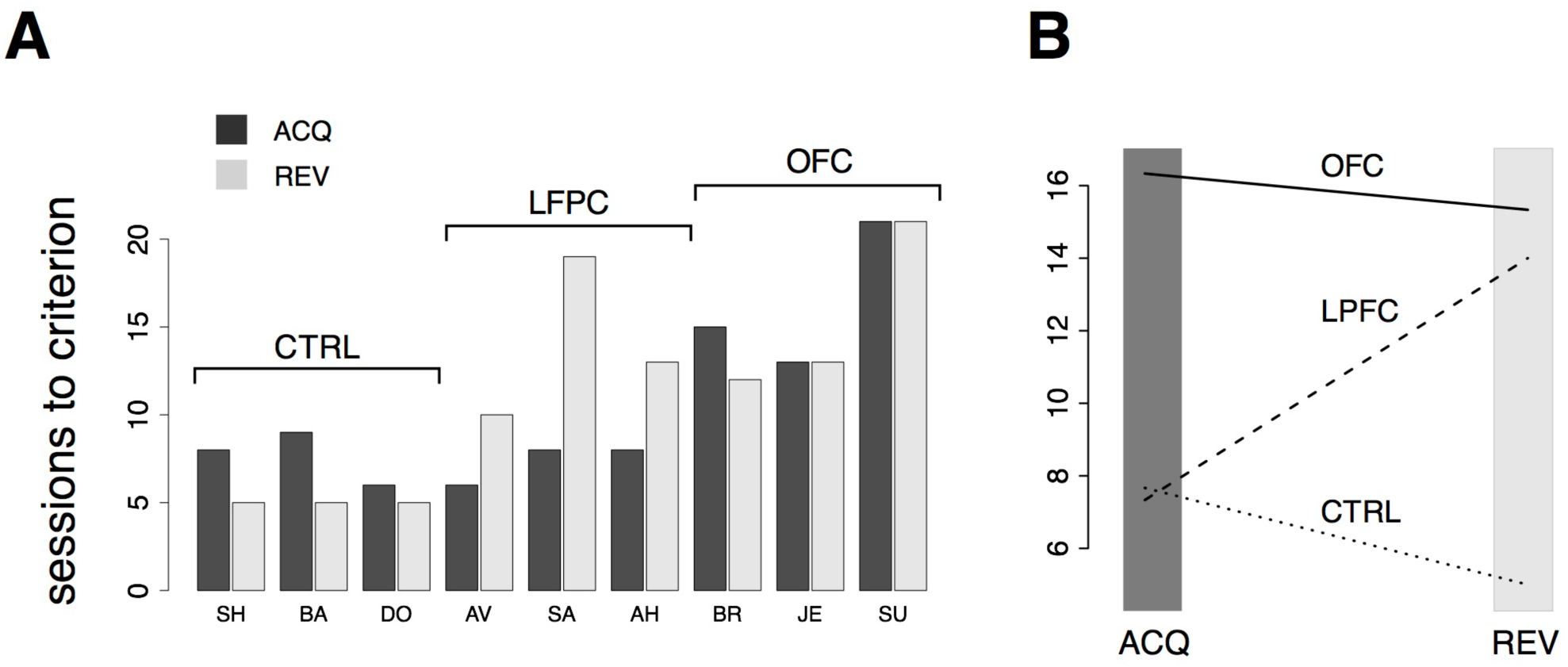
Analysis of behavior in the DMS task. **A)** Number of sessions-to-criterion for acquisition (“ACQ”) and reversal (“REV”) of the DMS task for control, LPFC, and OFC monkeys. A significant difference between groups was found (2-way, mixed design ANOVA, p<0.02). **B)** Interaction plot (protocol × group) of the 2-way ANOVA used to compare mean sessions-to-criterion among control, LPFC, and OFC monkeys shows an impairment of OFC monkeys in acquiring and reversing the task and an impairment of LPFC monkeys in reversing the task, whereas acquisition performance of LPFC monkeys was not significantly different than controls. See the text for details.

The rule-outcome association was then reversed (reversal marked by vertical lines in Figure 3A) between 2 and 5 sessions after acquisition (see Methods section “*DMS task”*). Performance dropped to ~50% correct in all monkeys by the end of the first session after reversal (Figure 3A; for monkey SU in the OFC group the overall score of the entire session was significantly different than 50%, but the 50% mark was reached by the 4^th^ quarter of the session; not shown). All 3 control monkeys acquired the reversed task within 5 testing sessions. However, this time both the LPFC and the OFC monkeys took significantly longer to reverse (LPFC monkeys: 10 to 19 sessions, Figure 3A, middle row; OFC monkeys: 12 to 21 sessions, Figure 3A, bottom row).

Recall that the monkeys could ‘skip’ a trial in one of two ways: either by releasing the bar too early (within 200ms from the onset of green), or by not releasing the bar at all (within 1 second after onset of green). The same percent correct performance can result from different patterns of ‘skip rates’.

For example, 50% correct could be the result of never skipping a trial as well as skipping all trials. In Figure 3B we show the skip rates in match vs. non-match trials, before and after reversal. As the monkeys learned the task, skip rates in the rewarded condition tended to decrease while the skip rates in the time-out condition tended to increase. Skip rates decreased to very low values in both conditions during the first session after a reversal, dropping the performance back to ~50% correct, as noted earlier. Despite some idiosyncratic features found in single monkeys (some tend to skip more and some less, in general), these trends were found in monkeys of all groups. Similarly, we found no clear difference in the pattern of ‘early’ (before green) vs. ‘late’ (>1s after green on) bar releases among different groups (not shown), suggesting a lack of clear difference in *how* the rules of the task were acquired by monkeys of different groups. When the monkeys started to skip time-out trials, they did so by releasing late, and then switched to releasing early after considerable practice (typically, after the control tasks, when exposed to the DMS task with short delay – see Methods for details on the sequence of tasks). This pattern was found in all monkeys.

The numbers of sessions to criterion are summarized in Figure 4A and were subjected to a 2-way, mixed design ANOVA. We found a significant difference among the groups (p<0.03) and a significant interaction of the effects of group and protocol (i.e., acquisition vs. reversal; p<0.01). Inspection of the interaction plot (Figure 4B) suggests that significant interaction is to be attributed to the LPFC group. Post-hoc comparisons confirmed that acquisition in this group was similar to controls (p>0.8, Mann-Whitney test) but reversal learning was significantly impaired (p=0.0318, Mann-Whitney test, one-tailed). During testing in (but prior to the acquisition of) the DMS task, the mean reaction times for the treated monkeys were not significantly different from those of control animals (2-way, mixed design ANOVA, no effect of group or group-trial type interaction, p>0.27), showing that the lesions caused no motor impairment.

### Dependence of performance on delay duration and novel stimuli

What strategy did the monkeys use to learn the DMS task? We sought to answer this question by analyzing how the performance depended on i) the temporal delay between the visual images or ii) the use of a new cue set. Regarding i), if the monkeys were relying on working memory to perform the task, their performance should degrade with lengthened delay duration, as previously reported (Fuster and Bauer, 1974; Bauer and Fuster, 1976; Shindy et al., 1994; Petrides, 2000a). As for ii), to confirm that the monkeys were using the concept of ‘same’ vs. ‘different’ rather than simply form hundreds of stimulus-outcome associations, we tested their ability to generalize the rule to a new cue set. Given that we used sets of 50 stimuli in each session, there were 2,500 possible combinations of matching and non-matching pairs, leading to the hypothesis that the monkeys used the information contained in the abstract concepts of ‘same’ and ‘different’, independent of cue identity. Based on this hypothesis, performance with a new cue set should not be different than the previously acquired level.

We tested both hypotheses concurrently by gradually increasing the delay up to 21 seconds while occasionally introducing a new cue set. Delay duration was increased by one second after each day of significant percent correct with the previous delay (see Methods for details). Performance was idiosyncratically related to delay duration, increasing with duration (and practice) for some of the monkeys and decreasing for others, regardless of group (Figure 5A). The differences in percent correct across groups were not significant (Kruskal-Wallis test, p>0.49). Thus, in our version of DMS, increasing the interval between the sample and test images did not necessarily lead to a decline of performance, independently of the manipulation (control, LPF or OFC lesion).

**Figure 5:**
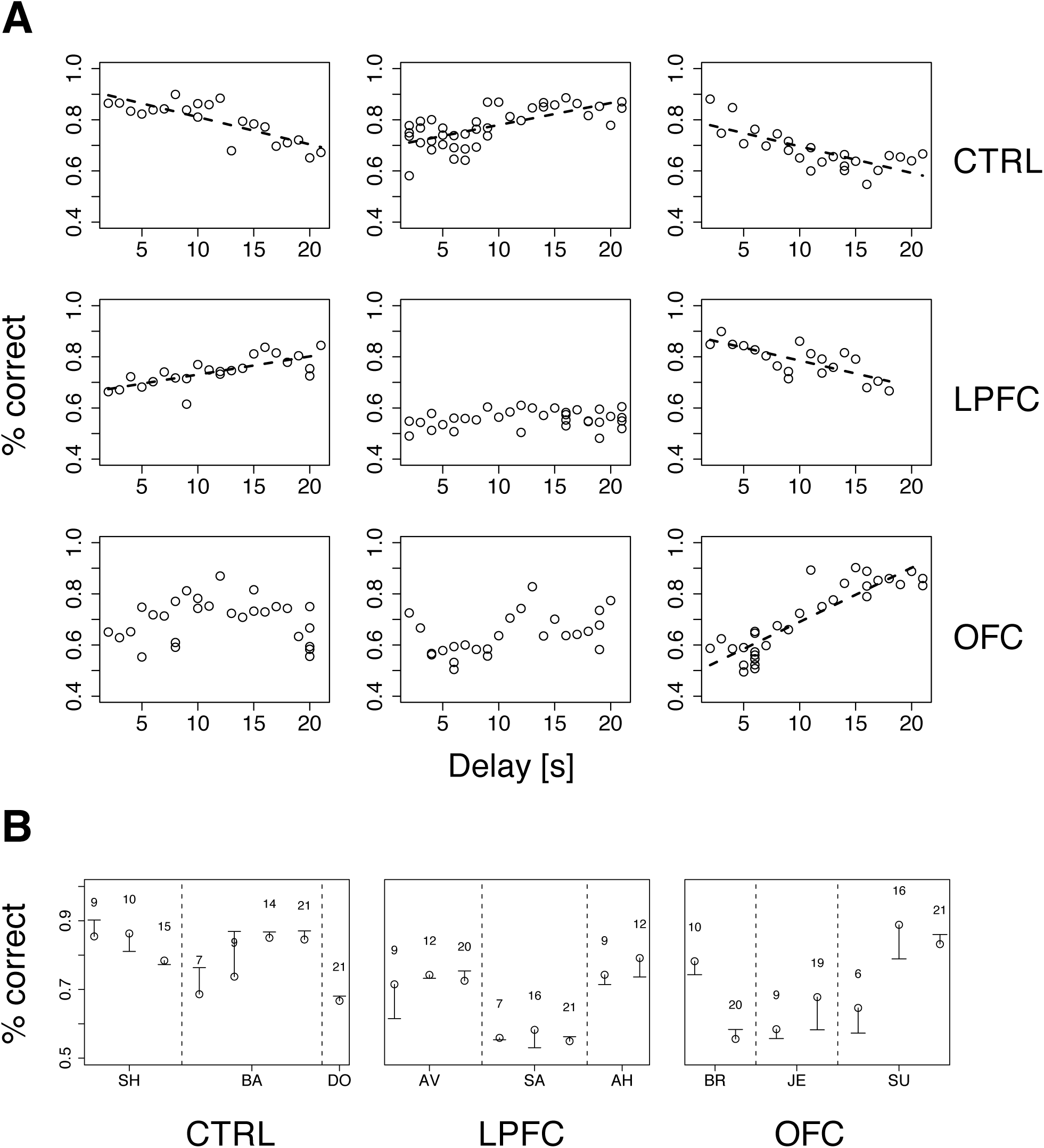
Performance vs. delay duration in the DMS task. **A)** Percent correct vs. delay duration for all monkeys (ordered as in Figure 3). Top, middle and bottom rows refer to control (CTRL), LPFC and OFC monkeys, respectively. When significant (p<0.01), regression lines are shown as broken lines (among all significant regressions it was p<0.0003). See Methods for details. **B)** Percent correct with familiar (circles) and new cue (dash) sets for all monkeys. The novel set was introduced one day after familiar one. The numbers above symbols indicate the temporal delay used for both sets (seconds). Vertical broken lines separate data from different monkeys within the same group. See Methods for details.

While increasing delay duration, new cue sets were occasionally introduced for each monkey, using the same delay as in the immediately preceding session (see Methods for details). The new cue set did not affect performance (Figure 5B), indicating that the monkeys were using the abstract concepts of ‘same’ and ‘different’ regardless of the individual identity of the visual stimuli. It is also clear, from Figure 5A, that all monkeys attained a similar maximum level of performance, albeit at different points during testing (at the beginning, at the end, or in the middle) within groups as well as across groups. In conclusion, the lesions affected acquisition and/or reversal learning rates (Figure 3), but not the asymptotic levels of performance or the dependence on delay duration. Moreover, the lesions slowed down, but did not abolish, the ability of the monkeys to form an abstract concept, because all monkeys performed well with a new cue set.

### The monkeys’ impairment did not depend on delay

Correct performance of the DMS task requires several steps: i) remembering a visual stimulus across a temporal delay ii) forming the ‘same’ vs. ‘different’ abstract rules, iii) learning to predict the outcome of each action depending on the rule in effect, and iv) adopting a suitable behavioral strategy to take advantage of this knowledge. In the previous section, we have investigated the roles of delay duration and the formation of an abstract rule on the monkey’s behavior. However, we have not investigated the role of the delay *per se* (as opposed to no delay at all) or the role of the abstract concept *per se* (as opposed to no concept at all). To understand if observed impairments were due to the presence of a temporal delay, and/or to the necessity to form the abstract concept of ‘same’ vs. ‘different’, we performed three control experiments, each requiring a subset of the abilities summarized above, but all having the same fundamental task structure (release on green to obtain a reward or a time-out).

In the first control, one of two new visual cues predicted a reward, while the second cue predicted a time-out (Figure 1E). Each cue was present during the entire duration of the trial. Everything else was as in the DMS task (see Methods for details). Since the monkeys learned to distinguish between the stimuli within 1 testing session, and also reversed the task in 1 session, we measured performance in terms of the number of trials to criterion, in hopes to resolve differences occurring within a single session. Trials-to-criterion were obtained with an iterative change-point procedure (Gallistel et al., 2001; Gallistel et al., 2004) (Figure 2 and Methods) and then analyzed with a 2-way, mixed design ANOVA (Figure 6A). There was a significant group effect (p=0.048), with the LPFC requiring more trials (93±68, mean ± SD) than either the control or the OFC group (44±26, mean ± SD over both groups). However, there was no difference in the rate of initial acquisition vs. the rate of reversal (acquisition vs. reversal, p>0.3), nor was there an interaction between rate of acquisition/reversal and group (p>0.9). Learning took less than 180 trials across all monkeys and was much faster than in the DMS task.

**Figure 6:**
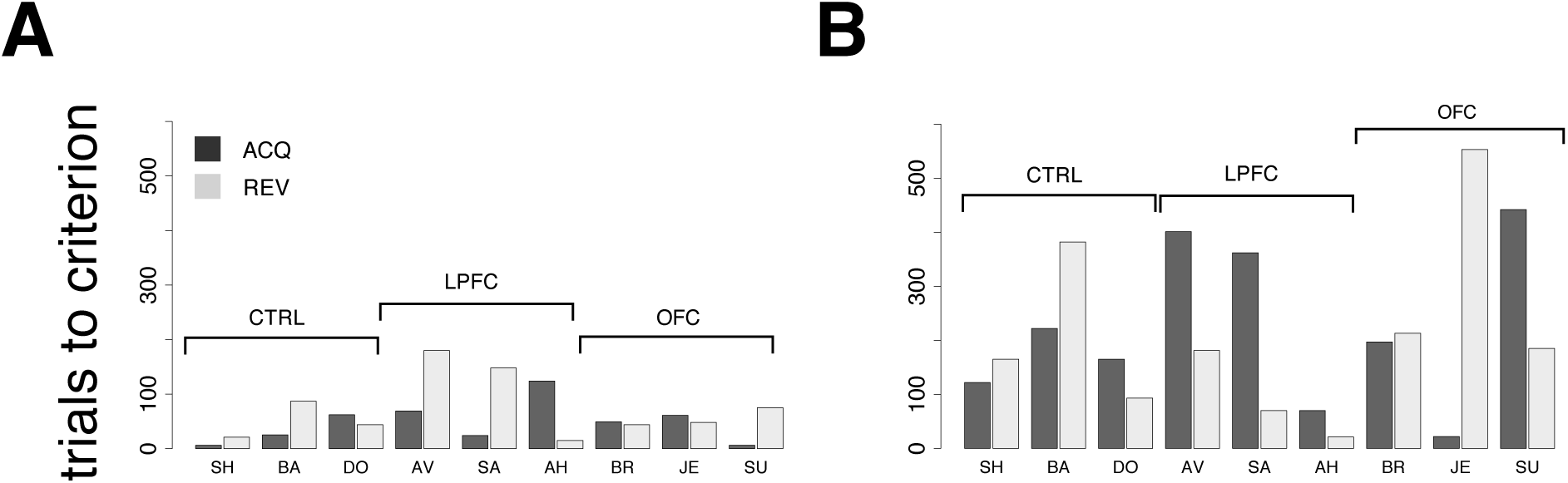
Behavior in control tasks. **A)** Number of trials to criterion for the cue-outcome association task of Figure 1E (all monkeys). Dark-grey bars: trials to acquisition; light-grey bars: trials to reversal. See the text for details. **B)** Trials to criterion for the delayed cue-outcome association task of Figure 1F (all monkeys; same key as in panel A).

Qualitatively, these results did not change when a delay was introduced between the cue and the response (Figure 1F). In this variation of the task, the cue or its meaning had to be remembered across the delay. As with the previous task, all monkeys acquired the delayed task within one testing session, and trials-to-criterion were computed as described above (Figure 6B). No significant effect of either group or protocol was found on the number of trials-to-criterion (p>0.5, 2-way ANOVA, mixed design; 215±150 trials-to-criterion, mean ± SD over all groups and both protocols). Learning was slower than in the simpler cue-outcome association task without delay, but it was still much faster than in the DMS task.

### The monkeys’ impairments relate to abstract concept formation

Finally, to investigate the role of the abstract concepts of ‘same’ vs. ’different’, we re-tested the monkeys in the DMS task. This time, the mnemonic requirements were kept to a minimum by using only a 100ms delay between sample and test (see Methods for details). The monkeys were tested in the same DMS task to which they had been last exposed (days-to-criterion shown as light bars in Figure 4A and reproduced in Figure 7 as white bars). All 9 monkeys performed well above chance on the first testing day, and then at least 3 sessions were needed to reverse the task (black bars in Figure 7): three sessions for controls, 4 to 11 sessions for lesioned monkeys, with a small but significant effect of group (p=0.049, Kruskal-Wallis rank sum test) but no difference between LPF and OFC monkeys (see Figure 7, black bars).

**Figure 7:**
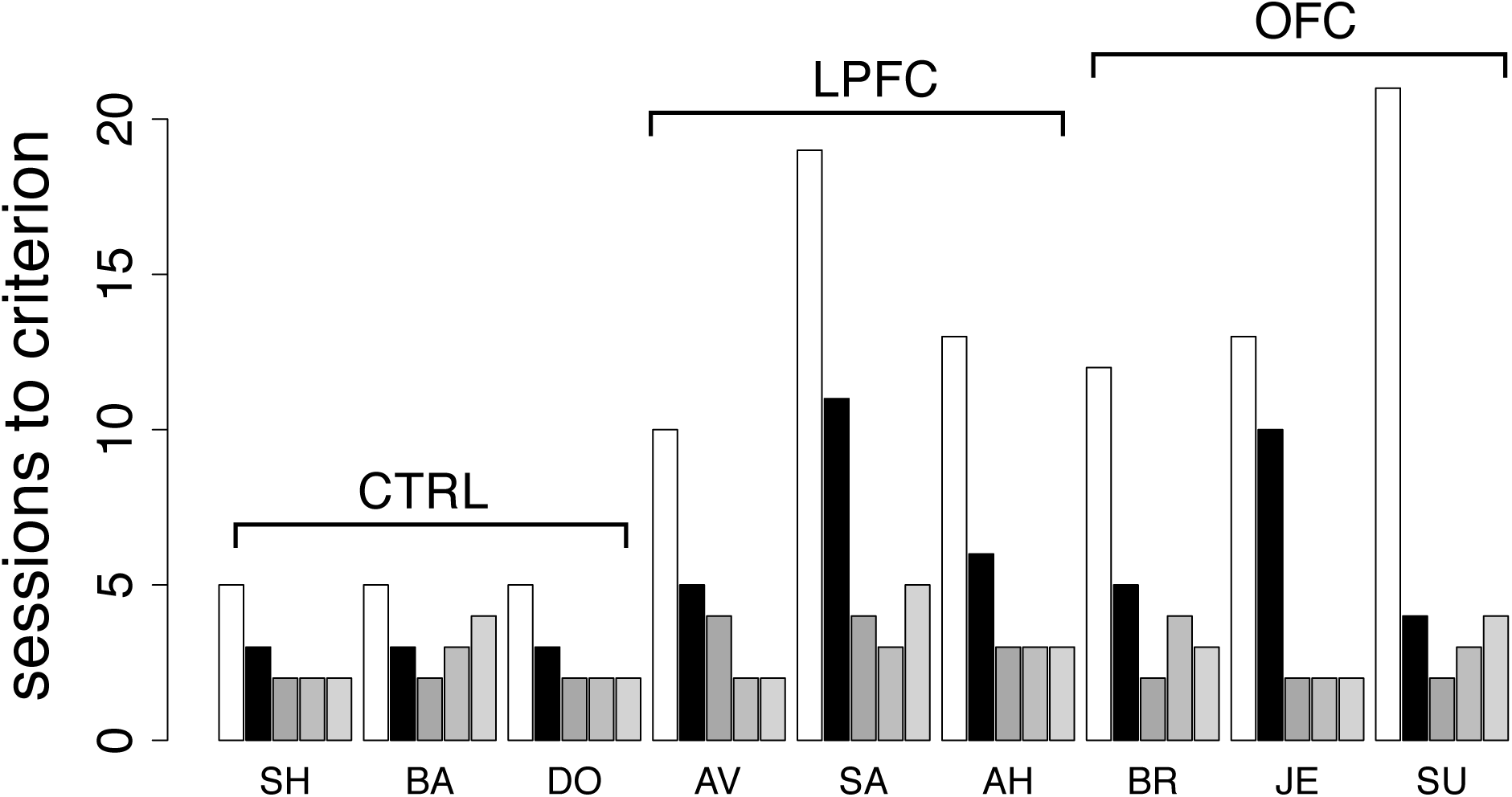
Behavior in the DMS task with 100ms delay and in additional reversals of the DMS task. Number of sessions required for reversing the DMS task with 100ms delay for all monkeys (black bars) and for 3 additional reversals of the DMS task with 1s delay, temporally ordered from left to right (grey bars). For comparison, the number of sessions required to reverse the DMS task with 1s delay are also reported (white bars, same as light grey bars in Figure 4A). Decrease in learning times with the number of reversals reveals the presence of a learning set. See the text for details.

Differently from the previous tasks based on cue-outcome association rules instead of rules based on abstract concepts, it took longer than one session to reverse the DMS task with 100ms delay, with a significant effect of either lesion. This suggests that the necessity to form an abstract concept to perform the task is key to the impairments. We also note, however, that learning to reverse the DMS task with 100ms delay was faster than learning to reverse the DMS with 1 second delay in all monkeys (Figure 7, compare black to white bars). Thus, the presence of a significant temporal delay could still be responsible for the impairments, at least to a partial extent. The alternative hypothesis is that the monkeys were developing a ‘learning set’, that is, they were learning about the reversals through repeated practice with them. To test this hypothesis, we increased the delay to 1 second and reversed the task three additional times. For all groups, performance kept improving over the three additional reversals (Figure 7, grey bars), and the difference among groups disappeared (2-way ANOVA, mixed design, p>0.28). This also shows that despite the difficulty with the initial rule reversals, the ability to form a learning set for reversal of the abstract rule was not impaired in LPFC and OFC monkeys, similar to what found in marmoset monkeys in a simpler association task (Dias et al., 1997).

Taken together, these results imply that the most taxing cognitive component in reversing the DMS task was learning the new association of the concepts of ‘same’ and ‘different’ with the reward-contingency, with a much smaller role (if any) for short-term memory (Rushworth et al., 1997) or for reversing a cue-outcome association per se.

## Discussion

Rule-based behavior is advantageous because no new learning is required when the parameters of a task change (here, the identity or the number of the images, and the length of the delay interval), allowing flexibility in mapping circumstances into actions. Here, we have shown that rhesus monkeys can rapidly learn to predict forthcoming behavioral outcomes using a rule based on the abstract concept of ‘same’ vs. ‘different’, and that their behavior in this task is differentially affected by bilateral removals of the lateral or orbital frontal cortex. Monkeys with bilateral OFC removals learned and reversed the rule significantly more slowly than controls. Perhaps more surprisingly, monkeys with large bilateral LPFC lesions learned the same task just as quickly as unoperated controls, but were significantly slower at carrying out the first rule reversal. Crucial to the impairments observed in lesioned animals was that the rule of the task was based on the abstract concepts of ‘same’ vs. ‘different’, since the lesions did not affect the ability to learn simpler direct cue-outcome associations that did not require the formation of an abstract concept.

It is thought that PFC is essential for rule learning (Burgess, 2000; Procyk et al., 2000; Bussey et al., 2001; Miller and Cohen, 2001; Wallis et al., 2001; Wood and Grafman, 2003; Genovesio et al., 2005; Tanji and Hoshi, 2008; Wise, 2008; Rigotti et al., 2013). The rapid learning rates and the lack of effect of LPFC lesions on acquisition of our DMS task seem at odds with previous results. It is unavoidable that different studies differ in the exact extent of the lesions, which is often mentioned as the potential reason for confounding and contradicting results in the literature. In our case, however, we had large LPFC lesions extending to both the superior and inferior convexity and including the principal sulcus, with little opportunity for remaining fragments of LPFC tissue. It has been reported that such extended lesions impair performance in matching tasks in monkeys with or without a spatial component (Passingham, 1975; Mishkin and Manning, 1978), and with (Fuster and Alexander, 1971; Goldman-Rakic, 1995, 1996; Miller et al., 1996; Petrides, 2000b; Curtis and D'Esposito, 2003, 2004), or without (Rushworth et al., 1997), a delay. Thus, we expected that our LPFC monkeys would be impaired in the acquisition of the DMS task. Instead, we found that the LPFC monkeys were as facile as controls in acquiring the task, and were instead impaired at reversing it. This deficit does not seem to be of a perseverative nature (Dias et al., 1997; Clarke et al., 2008; Buckley et al., 2009), since the monkeys quickly abolished their previously acquired strategies (i.e., release on green in rewarded trials only), and started releasing on green in all trials after the task rule was reversed (cf. Figure 3), presumably to sample as many rule-outcome contingencies as possible.

If the exact extent of the LPFC lesions is not to blame for the lack of impairment in acquiring the DMS task, how can one explain the difference with previous findings? We believe that the specific structure of the tasks used here is the most likely explanation. Unlike procedures involving touching computer screens, or direct manual object displacements as in the Wisconsin General Testing Apparatus (WGTA, see e.g. (Moore et al., 2009, 2012) for recent studies), learning DMS in an automated apparatus can be laborious for monkeys, sometimes taking months (Pigarev et al., 2001); yet, our monkeys displayed rapid learning times. This suggests that the monkeys would learn to perform our tasks in a different way compared to other studies. All previous studies, for example, involved an active action selection process, i.e., different motor actions in different conditions (such as match vs. non-match trials in the DMS task). By removing the action selection process altogether, we stripped the task of any differential motor component. Although the monkeys eventually learned to produce the bar release at different times to complete or ‘skip’ trials, they were initially required to perform only one action: to release the bar on green. Once they had learned to release on green and sample the outcome in each trial type, they could act on their knowledge and intentionally ‘skip’ unrewarded trials. The behavior of our control monkeys alone proves that this strategy is fundamentally different from learning, simultaneously, the meaning of trial types and a desired response, eliminating a perennial confounding aspect of these types of tasks and resulting in much faster learning times.

Two factors of our procedures seem most important in producing this result. First, the monkeys are exposed to the DMS task only after having been taught to release a bar after a red target turned green. The initial training in the bar release task, which is easily learned (~2 weeks, see e.g. (Liu et al., 2004)), taught the monkeys about the only operant action required to perform our DMS task. Second, when testing the monkeys in the DMS task, we capitalized on the tendency of the monkeys to quickly learn the meaning of predictive cues and to abort unrewarded trials prematurely. As we know from previous studies (Bowman et al., 1996; La Camera and Richmond, 2008; Minamimoto et al., 2009), incurring a delay before receiving a reward is a powerful modulator of the monkeys’ behavior, that often results in incorrect or premature bar release even when correction trials are required. In our DMS task, learning this behavior is further facilitated by the fact that incorrectly timed bar release results in an *advantageous* strategy that the monkeys learned quickly. With different variations of this procedure, we have found that monkeys also learn attentional sets and categories quickly (Lerchner et al., 2007; Minamimoto et al., 2010). This design seems to tap into the monkeys’ natural ability to infer information from the environment (Bromberg-Martin and Hikosaka, 2009) and take advantage of it, somewhat similarly to what has been observed with the repeat-stay, change-shift strategy (Bussey et al., 2001).

The other key aspect of this study is the direct comparison in tasks that differ only for the presence of a DMS component. This allowed us to disentangle the case where it is an abstract concept (‘same’ vs. ‘different’) that is predictive of reward contingency, from the case where it is a single visual stimulus. In addition, since the concepts of ‘same’ vs. ‘different’ had to be formed across a temporal delay (up to 21 seconds), we could explicitly test the importance of the delay between cue presentation and motor response. Our main finding is that only when the abstract concept is present we detect a clear and statistical significant impairment of the lesioned monkeys (Fig. 6B). On the one hand, the results on acquisition are in keeping with previous studies finding a largely intact ability of monkeys with OFC damage to make appropriate choices when initially learning the values of available options (Izquierdo et al., 2004; Clarke et al., 2008; Walton et al., 2010). On the other hand, these results are at odds with the known involvement of OFC in reversal learning guided by conditioned stimuli, which is typically accompanied by pathological perseveration (Murray et al., 2007; Clarke et al., 2008). Once again, the most likely explanation is due to the difference in task procedures, which in our case involve a single action that had been previously learned. It is also possible that, having learned the more taxing DMS task first, our lesioned monkeys found easier to learn the simpler control tasks. Additional experiments are required to resolve this ambiguity – here, we had to test the monkeys in the DMS task first, to make sure that acquisition of DMS would not be facilitated by previous testing in the simpler cue-outcome association tasks. The fact that acquisition of our DMS task was impaired in OFC monkeys, whereas reversal of the simpler cue-outcome associations were not, reveals that OFC is essential for forming an abstract concept and/or for correctly assigning a predicted outcome to it (more work is necessary to establish which).

In summary, by comparing the effect of LPFC and OFC lesions in different scenarios, one of which requires forming an abstract concept, we found that both LPFC and OFC are essential for different aspects of learning the association between an abstract concept, a decision, and a predicted outcome. Neither of these brain regions seemed crucial in establishing a learning set for concept-outcome associations. It appears that OFC is needed to form the “same/different” concepts initially, learn the rule and/or associate it with the outcome, whereas LPFC seems to be needed to modify a previously acquired rule-outcome association.

## Acknowledgments

We are indebted to R.C. Saunders and E.A. Murray for performing the lesions. We thank D.P. Soucy and J. Peck for their technical assistance, T. Minamimoto, A. Lerchner, J.M. Simmons, S. Ravel, M. Mishkin and R.C. Saunders for useful discussions. This study was supported by the Intramural Research Program of the National Institute of Mental Health. During the completion of this work G.L.C. was supported by NIH ARRA P30 NS069256 start up funds. The views expressed in this article do not necessarily represent the views of the NIMH, NIH, or the United States Government. The authors declare no competing financial interests.

